# Experimental bleaching of photosymbiotic amoeba revealed strain-dependent differences in algal symbiosis ability

**DOI:** 10.1101/2024.07.24.604942

**Authors:** Daisuke Yamagishi, Ryo Onuma, Sachihiro Matsunaga, Shin-ya Miyagishima, Shinichiro Maruyama

**Author notes:** Corresponding authors: Shin-ya Miyagishima, Shinichiro Maruyama.

## Abstract

Photosymbioses, the symbiotic relationships between photosynthetic algal symbionts and non-photosynthetic eukaryotic hosts, are sporadically found in a lot of eukaryotic lineages, but only a few taxa, such as cnidarians and ciliates hosting algal endosymbionts, have been actively studied for a long time. That has hindered understanding the universal mechanisms of the photosymbiosis establishment. Especially in Amoebozoa, only two species, *Mayorella viridis* and *Parachaos zoochlorella*, are reported as photosymbiotic in nature, and their mechanisms of establishing symbiotic relationships are still unclear. To investigate the extent to which and how photosymbiotic amoebae depend on the symbiotic relationships, *M. viridis* were treated with reagents that are known to induce the collapsing of photosymbiotic relationships, or bleaching, in other photosymbiotic species. As a result, we succeeded in artificially removing algal symbionts from host *M. viridis* cells with an herbicide, 2-amino-3-chloro-1,4-naphthoquinone. The apo-symbiotic state amoeba cells were able to survive and grow to the same extent as the symbiotic state cells when they fed microbial prey, indicating that the algal symbionts are not essential for the host growth under certain conditions. Furthermore, to see whether the photosymbiotic state is reversible, we fed two strains of algal symbionts to the apo-symbiotic amoeba host. The result showed that the apo-symbiotic hosts were able to ingest symbiont cells and re-establish the symbiotic state. The efficiencies of ingesting algal cells were significantly different depending on algal symbiont strains, indicating that different algal strains possess discrete symbiotic abilities to *M. viridis*. To our knowledge, we provide first insights on the establishment and collapse of photosymbiosis in Amoebozoa, which pave the way to understand the universal mechanism of photosymbiosis utilizing *M. viridis* as a model system.

## Introductions

Aquatic photosynthesis by algae and photosynthetic prokaryotes is responsible for approximately half of net-productions on the earth (Field *et al*. 1998). Some microalgae, as symbionts, can establish symbiotic relationships with non-photosynthetic eukaryotes as hosts, which is called photosymbiosis. Photosymbiosis has been reported from various aquatic and terrestrial environments. For example, corals, sea anemones, and other cnidarians belonging to Opisthokonta are known to have photosymbiotic relationships with some microalgae including dinoflagellates and green algae. Photosymbioses between ciliates and *Chlorella* have been also well studied. Some foraminifera species, which are members of Rhizaria, have photosymbiosis relationships with several lineages of algal symbionts, such as dinoflagellates, red algae, diatoms (Johnson 2011, Dorrell and Howe 2012).

The mechanisms of photosymbioses are studied from a molecular, physiological, and ecological perspective particularly in cnidarians and ciliates (Davy *et al*. 2012, Fujishima and Kodama 2012). Importantly, researchers can artificially “bleach”, or induce the apo-symbiotic state, in which all the algal symbionts are expelled from their host bodies, of these species, which is one of the reasons that make them model organisms. There are several reports about artificial bleaching methods so far (Belda-Baillie *et al*. 2002, Mihirogi *et al*. 2003). For example, the model sea anemone Aiptasia (*Exaiptasia diaphana*) can be bleached by exposure to a high temperature of 35°C for 20 days (Belda-Baillie *et al*. 2002), or more easily by treating them with a herbicide 2-amino-3-chloro-1,4-naphthoquinone (ACN), which is also known as quinoclamine (Mihirogi *et al*. 2023). Green hydra also belonging to *Cnidaria* and having green algal endosymbionts can be bleached as well by treating with glycerol (Whitney 1907). Similarly, some bleaching induction methods have been developed for ciliates. *Paramecium bursaria* intracellularly possessing *Chlorella* endosymbionts can be bleached by treatment with paraquat for 5 days (Hosoya *et al*. 1995). There are reports that cycloheximide and 3-(3,4-dichlorophenyl)-1,1-dimethylurea (DCMU) are also effective in bleaching *P. bursaria* (Kodama and Fujishima 2008, Reisser 1976). Using these bleaching techniques, we can compare and analyze the responses when the hosts and symbionts are in a symbiotic state or apo-symbiotic state. In the case of the sea anemone *E. diaphana*, the changes in metabolisms of the hosts in response to symbionts can be studied by comparing transcriptomes of the symbiotic state with the apo-symbiotic state (Lehnert *et al*. 2014, Ishii *et al*. 2019). Similar studies have been conducted on green hydras and *P. bursaria* (Ishikawa *et al*. 2016, He *et al*. 2019).

Although eukaryotic species possessing photosymbionts have been found sporadically within phylogenetically distant taxa, such as Opisthokonta and Alveolata, and a limited number of model species have been studied. Thus, it is still unclear whether cellular or molecular features found in some photosymbiotic organisms are universal or lineage-specific. Especially among a lot of Amoebozoans that have predatory behavior suitable for observing the phagocytosis process, which has been known as the first and key step of photosymbiosis establishment (Davy *et al*. 2012), only two amoebozoan species are found as photosymbiotic organisms and extremely few studies have been conducted in their photosymbiosis establishments at molecular and cellular levels. These have hampered understanding of the importance of the phagocytosis process in photosymbioses, and it is important to build an ideal system in which we can simply compare the predatory process and symbiosis process.

*M. viridis* is a member of Amoebozoa in freshwater and intracellularly possesses microalgal endosymbionts, *Chlorella* sp. (Cann 1981). Because the amoeba cells are adhesive and their locomotion is slower than some other photosymbiotic protists, such as the ciliate *P. bursaria*, it is comparatively easy to observe the algal symbionts inside the amoeba host cells under microscopy. Taking advantages of this species, here we show that we have developed a bleaching method for the amoebae and made a quantitative comparison between different *Chlorella* strains.

## Materials and methods

### Cell culturing

*M. viridis* (strain name og-1, provided by Iwakuni city micro-life museum, Japan, originally isolated and cultivated by Dr. Masashi Hayakawa at Osaka University) was cultured in 8ml of sterilized mineral water “OISHIIMIZU ROKKO” (hereinafter called RW, Asahi Soft Drinks Co., Ltd., Japan) in Cell Cultivation Flask (Vent Cap) 25 cm^2^ (VTC-F25V, AS ONE CORPORATION, Japan). Cultivation light and temperature were 8 μmol s^-1^ m^-2^ constant light and 20°C respectively. The medium in the flask was changed when the base of the flask got dirty. First, the medium was discarded and 8 mL of fresh RW was newly added. The flask was gently rinsed with the added medium, the medium was discarded again, and 8 mL of fresh RW was added. Then, the bottom of the flask was tapped about 20 times to suspend the amoebae cells and immediately the culture medium was transferred into a new cultivation flask.

*Cryptomonas paramecium* (NIES-715) was used as feed for *M. viridis. C. paramecium* was maintained in 100 ml CYT medium (https://mcc.nies.go.jp/medium/ja/cyt.pdf) in Cell Cultivation Flask (Vent Cap) 75 cm^2^ (VTC-F75V, AS ONE CORPORATION, Japan). Cultivation light and temperature were approximately 40 μmols^-1^s^-2^ 12L:12D and 20°C respectively. For one *M. viridis* culture flask, 10 ml of *C. paramecium* culture for was used. The culture was centrifuged at 1000 × *g* for 10 minutes and the supernatant was removed. Ten ml of sterilized RW was added to the pellet and the pellet was suspended by pipetting. The suspension was centrifuged at 1000 × *g* for 10 min again and the supernatant was removed. Finally, 1mL of sterilized RW was added to the pellet to resuspend by pipetting, and added to each *M. viridis* culture flask.

Two *Chlorella* strains (strain A and B) were isolated from *M. viridis* culture and grown in 20 mL AF-6 medium in a Cell Cultivation Flask (Vent Cap) 25 cm^2^ (VTC-F25V, AS ONE CORPORATION, Japan) at 20°C under approximately 37 μmol s^-1^ m^-2^ 12L:12D.

### Screening for bleaching reagents

Cultivation flasks of *M. viridis* were tapped about 20 times to suspend the amoebae cells and the 1 ml of cell suspension was added to every 5 random wells of a 48-well plate (VTC-P48, AS ONE CORPORATION, Japan). To test bleaching reagents, 1μl each of 70% ethanol, sterilized RW, 5 mM ACN (FUJIFILM WAKO Pure Chemical Corporation, Japan) solution in 70% ethanol (final concentration: 5 μM), and 5 or 50 mM DCMU (FUJIFILM WAKO Pure Chemical Corporation, Japan) solution in 70% ethanol (final 5 or 50 μM, respectively) were added into each well. The plate was maintained at 20°C under 8 μmol s^-1^ m^-2^ constant light for one week. The experiment was repeated three times.

The amoebae cells on the bottom of the plates were suspended by pipetting and the 50μl cell suspension was taken from the wells and dropped onto a glass slide (S1214, MATSUNAMI GLASS IND., LTD., Japan). This manipulation was repeated until an amoeba cell was found in the droplet under 20x brightfield microscopy observation (OLYMPUS CKX53, LCAch N). After confirming the amoebae were in the droplets, 18 × 18 mm cover glass (C218181, MATSUNAMI GLASS IND., LTD., Japan) was applied. The excess medium was removed with Kimwipes (Nippon Paper Cresia, Japan) until the amoebae cells of interest contacted both the glass slide and cover glass and stopped moving. Then, brightfield and autofluorescence images (CANON DS126761, OLYMPUS CKX53 Blue Excitation mirror unit) of the cells were acquired. The well samples in which amoeba cells were lost during the observation operation were not used for further analyses.

### Preparation of bleached *M. viridis* cultures

Eight ml of *M. viridis* culture was treated with 8 μL of 1 mg /ml ACN solution in 70% ethanol and incubated at 20°C under 8μmols^-1^m^-2^ constant light for one week. After the treatment, the medium of the culture was removed and 8ml of fresh sterilized RW was added to the culture for washing the cells. This manipulation was repeated twice. Washed bleached *M. viridis* cells were maintained at 20°C under constant dark to make them in the apo-symbiotic state. The methods of feeding and medium change were the same as *M. viridis* cells in the symbiotic state.

### Measurement of growth of the *M. viridis*

Before each measurement of the cell number, the *M. viridis* culture was washed by removing the medium with an aspirator, adding 8ml of fresh RW, and shaking the flask lightly. The fresh medium was removed again and 8ml of fresh RW was added. 100 μL of the washed *M. viridis* cell culture was transferred into a randomly selected well of a 96-well glass base well plate (GP96000, MATSUNAMI GLASS IND., LTD., Japan). 100 μL of *C. paramecium* cell culture washed with RW was added to the well so that the cell density of *C. paramecium* in the well was 2 × 10^6^ cells/mL. The sample plate was maintained at 20°C under 8 μmol s^-1^ m^-2^ 12L:12D. After tapping the plate to sink the amoebae cells to the bottom of the plate, the number of the amoebae in each well was counted under 20x brightfield microscopy observation (OLYMPUS CKX53, LCAch N). The amoebae in the symbiotic and apo-symbiotic states were used for this measurement (*n* = 4).

### Co-culturing of *M. viridis* and *Chlorella*

*Chlorella* culture was centrifuged at 1000 × *g* for 20 minutes. After removing the supernatant, the pellet was resuspended in an equal amount of sterilized RW with the *Chlorella* culture. This manipulation was performed twice to wash the *Chlorella* cells. Two wells of a 96-well plate (3870-096, AGC TECHNO GLASS Co., Ltd., Japan) were randomly selected. *Chlorella* cells were mixed into bleached amoebae cultures so that the algal density in the mixture was adjusted to 1.0 × 10^7^ cells/ml, to make 25 μL of cell suspension in each well. The mixture was incubated for 1 h.

### Quantification of *Chlorella* cells inside *M. viridis*

The amoebae cells were put off from the bottoms of the wells to resuspend by pipetting and a drop of the resuspension was taken from the well onto a glass slide. The observation method under microscopy was the same as the method described above. ROIs were created by surrounding amoeba cells in brightfield images with Polygon selections in Fiji (Schindelin et al. 2012). The ROIs were applied to the fluorescence image. Binarization of the fluorescence images and segmentation of the contacting the *Chlorella* cells were conducted by the Interactive H_watershed function of the Fiji plugin SCF-MPI-CBG (Seed dynamics = 1, Intensity threshold = 13, Peak flooding = 100). The number of *Chlorella* cells in the ROI was counted with Fiji’s ‘Analyze particles’ function. The average number of Chlorella cells in amoeba cells in the two conditions was tested by Welch’s t-test with R (R version 4.3.1, R Core Team (2023). _R: A Language and Environment for Statistical Computing_. R Foundation for Statistical Computing, Vienna, Austria. <https://www.R-project.org/>.).

### Phylogenetic analysis

Partial sequences of the small subunit (SSU) and internal transcribed spacer (ITS) of ribosomal DNA (rDNA) used for the phylogenetic analysis of *Chlorellaceae* were obtained from a previous study (Luo et al. 2010). In addition, PCR amplification of SSU and ITS of strain A and B were conducted with the following primers: primer set1, SR1_inf 5’-AGGTCGACTCTAGAGTACCTGGTTGATCCTGCCAG-3’ and SR12cF_inf_rev 5’-TTACGACTTCTCCTTCCTCTACTCTAGAGTCGACCT-3’ (modified Nakayama *et al*. 1996), and primer set2, SR12cF_inf 5’-AGGTCGACTCTAGAGTAGAGGAAGGAGAAGTCGTAA-3 and 25F1R_inf 5’-CGGTACCCGGGGATCATATGCTTAAATTCAGCGG -3’ (modified Takano & Horiguchi 2006). All sequences were aligned using MAFFT (ver. 7.487) with -linsi and -- adjustdirection options (Katoh & Standley 2013). The output file of the alignment was trimmed using TrimAl (ver. 1.4.1) with -gt 0.8 option (Capella-Gutiérrez 2009). A phylogenetic tree was created with the trimmed alignment file using IQ-TREE (ver. 1.6.12) with -alrt 1000 and -bb 1000 option (Nguyen et al. 2015, Kalyaanamoorthy et al. 2017).

## Results

### ACN can induce bleaching of *M. viridis* cells

To test the effects of potential bleaching reagents, we confirmed the presence or absence of algal symbiont cells inside the amoebae cells (i.e. the symbiotic or apo-symbiotic states) by observing chlorophyll autofluorescence of the symbionts when they were treated with a photosynthesis inhibitor DCMU and a herbicide ACN (Fig. 1 A). Chlorophyll fluorescence derived from symbiotic algae was observed in the amoeba hosts after one week of treatment in the control group with the mineral water RW and 70% ethanol, and the DCMU-treated cells. On the other hand, no chlorophyll fluorescence was observed in *M. viridis* cells treated with ACN for one week (Fig. 1 A). No abnormalities in cell shape were observed in bleached ACN-treated amoeba cells that had been after the ACN treatment and washing with RW (Fig. 1 B). These results indicate that ACN treatment can induce bleaching of *M. viridis* cells.

**Fig. 1.**
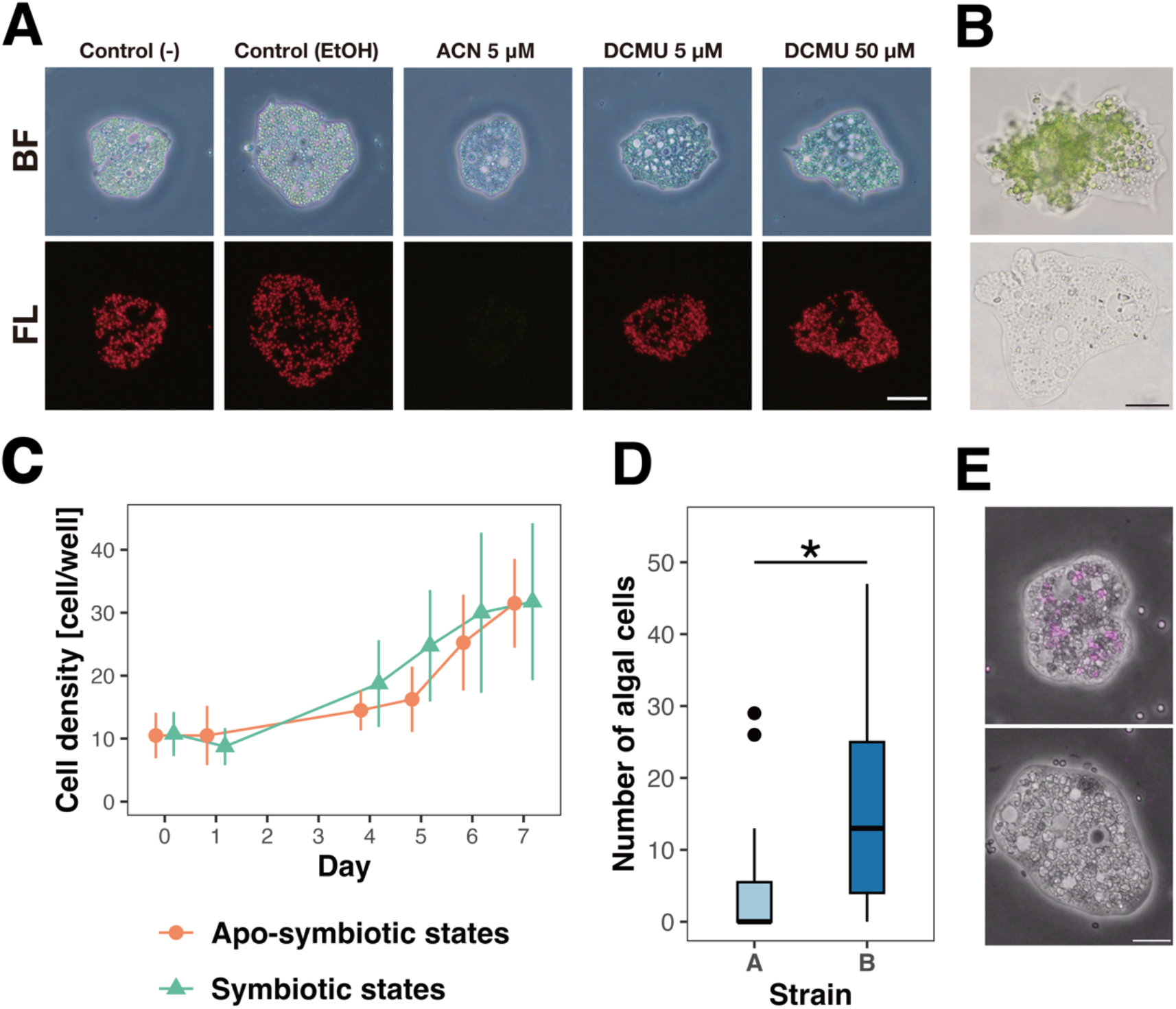
Bleached *M. viridis* and its cell biology. (A) Microscopic images of *M. viridis* in the bleaching experiment. After 1 week of treatments with no chemicals (-), chemical solvent (EtOH), ACN (5 µM), DCMU (5 µM, 50 µM), the amoeba cells in bright field (BF) and chlorophyll auto-fluorescence (FL) were observed. Scale bar = 40 µm. (B) Microscopic images of *M. viridis* in the symbiotic states (upper panel) and apo-symbiotic (bleached) states (lower panel). Scale bar = 10*μ*m. (C) The growth of *M. viridis* in symbiotic (SYM) and apo-symbiotic (APO) states. (D) Quantification of the number of cells of the *Chlorella* strains A and B ingested by *M. viridis*. The box represents the quartile numbers. The upper end and lower end of the bar represent the maximum value and the minimum value respectively. The plots represent the outliers. The asterisk indicates the significant difference (Welch’s *t*-test, *p*-value = 0.0242 < 0.05). (E) Examples of microscopic images of *M. viridis* co-cultured with *Chlorella* strains A (upper panel) and B (lower panel) for one hour. Bright field images are overlayed with chlorophyll auto-fluorescence of *Chlorella* cells. Scale bar = 20*μ*m.

**Fig. 2.**
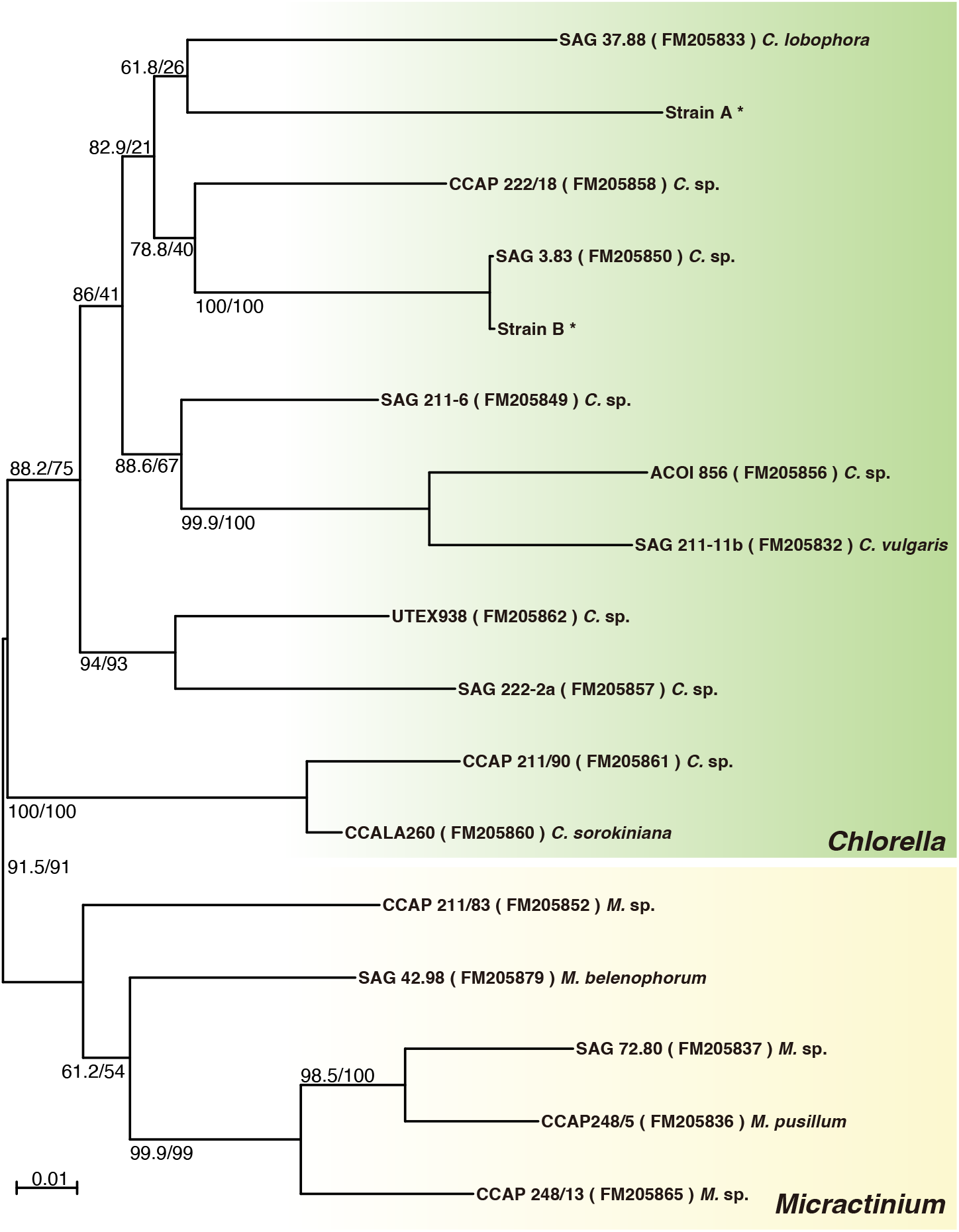
A phylogenetic tree of ITS and SSU rDNA regions in the genus *Chlorella*. The genus *Micractinium* was used as an outgroup. Each OTU is labeled with the strain name, the accession number of nucleotide sequence, and the species name, except for the ones determined in this study shown with asterisks (*). The values on each branch represent statistical support values, SH-aLRT (left) and UFboot (right).

### *M. viridis* can grow in the symbiotic and apo-symbiotic states

To verify whether and how the state of symbiosis could affect the *M. viridis* cells’ growth and survival in the apo-symbiotic state, we checked the growth of the amoebae in the symbiotic or apo-symbiotic state. The amoebae were cultured under constant light conditions and were fed *C. paramecium* for one week. The results showed that *M. viridis* in the apo-symbiotic state could grow to the same extent as the amoebae (Fig. 1 C). This indicates that symbiotic algae are not essential to survive and grow for *M. viridis* under certain conditions.

### Quantification of symbiotic abilities of algae to *M. viridis*

To confirm if *M. viridis* could re-establish the symbiosis, i.e. make a transition from the apo-symbiotic state to the symbiotic state, *Chlorella* cells were added into the apo-symbiotic *M. viridis* cultures. Two *Chlorella* strains isolated from *M. viridis*, strains A and B, were used to compare their symbiotic abilities. Bleached *M. viridis* cells were co-cultured with an excessive number of *Chlorella* cells for one hour. After the co-culturing, the number of *Chlorella* cells in the amoebae cells was counted. We found that the number of *Chlorella* cells ingested by *M. viridis* was significantly higher in strains B than A (Fig. 1 D). This result indicates that *M. viridis* in the apo-symbiotic state is able to re-establish symbiosis and retain the symbiotic state. In addition, the symbiotic abilities of two *Chlorella* strains are different, though these strains were isolated from the original *M. viridis* culture.

### Phylogenetic analysis of *Chlorella* strains

We conducted a phylogenetic analysis of the two *Chlorella* strains showing different symbiotic abilities, by using their ITS and SSU rDNA region sequences. The phylogenetic tree showed that both strains were in the genus *Chlorella*. Strain B was genetically close to the green algal endosymbiont (zoochlorella) in *Acanthochystis turfacea*, a member of Heliozoa. Considering that the strain B showed high symbiotic ability to *M. viridis* (Fig. 1D), the photosymbiotic ability of *Chlorella* may be maintained across phylogenetically distant host species.

## Discussions

### How photosymbiosis stability can affect the amoeba host

The establishment of photosymbiosis between *M. viridis* and algal symbionts does not seem to be correlated to the photosynthesis activity of the symbionts because DCMU suppressed the photosynthesis activity of free-living symbionts (see supplementary information) but did not affect the photosymbiosis stability (Fig. 1A). A previous study suggested that the symbiosis establishment between Aiptasia and symbiotic algae is independent of the symbiont’s photosynthesis (Jinkerson *et al*. 2022). Thus, the bleaching of *M. viridis* induced by ACN seems to be caused by factors other than the inhibition of the symbiont’s photosynthesis. Considering that ACN also induces the bleaching of cnidarians (Mihirogi *et al*. 2023), the same mechanism likely causes the symbiotic collapse in *M. viridis* as in cnidarians. Further studies on the molecular mechanisms by which ACN affects photosynthetic symbionts and host organisms will shed more light on how photosymbiosis is established.

Our result showed that the symbionts were dispensable for the survival and growth of *M. viridis* in our experiment setting. This leads to an idea that one of the reasons for the lack of reports of *M. viridis* in the apo-symbiotic state in field environments may be simply because the presence of symbionts is one of the characteristics used for species identification. Nevertheless, this does not necessarily mean that symbionts have no effect on the fitness of *M. viridis*. A previous study showed that the growth rates in the symbiotic and apo-symbiotic host ciliates were dependent on light and feeding conditions (Lowe *et al*. 2016). Meanwhile, the growth experiment in our study was conducted in a specific condition (constant light and excessive food). The balance between cost (symbiotic load) and benefits derived from the symbionts may fluctuate, e.g. depending on light and feeding conditions as shown in ciliates (Lowe *et al*. 2016), potentially leading to differences in the fitness of *M. viridis* between the symbiotic and apo-symbiotic states.

### Symbiotic ability of algal symbionts and symbiont recognition by host cells

The difference in symbiotic ability between the two *Chlorella* strains suggests that there is a symbiont recognition mechanism in *M. viridis*. Partner specificity in photosymbiosis has also been found in cnidarians (LaJeunesse *et al*. 2018, Yorifuji *et al*. 2021). A previous study on coral photosymbiosis suggests that the host corals can recognize their symbionts and modulate their immune system (Jacobovitz *et al*. 2021). Recent studies on other corals suggest that LePin (lectin and kazal protease inhibitor domain) proteins in hosts may be involved in phagocytosis and recognition of the algal symbionts (Hu et al. 2020, Hu et al. 2023). Lectins and sugar residues on the algal cell walls recognized by them may be a key for *M. viridis* to distinguish between symbionts and other organisms.

It is shown that amoebozoan predators that feed photosynthetic prey change the frequency of their phagocytic uptake and speed of food digestion depending on light conditions (Uzuka *et al*. 2019). Regulation of the balance between uptake, digestion, and expulsion of the algae by the host may have affected the difference in symbiotic abilities between the *Chlorella* strains used in this study. Comparative analysis of phagosome maturation of the apo-symbiotic *M. viridis* cells ingesting prey and symbiont may provide insights into this, and some hints to address evolutionary issues e.g. transitions from heterotrophic lifestyles to mixotrophic lifestyles in initial processes of plastid acquisition through endosymbiosis.

Strain B with high symbiotic ability to *M. viridis* was closely related to algal symbionts of *A. turfacea*, suggesting that the photosymbiotic ability of *Chlorella* species may be universal across phylogenetically distant host species. Quantification of the symbiotic abilities of strain B to other host species may shed light on this issue. Genomic analysis and development of genetic engineering methods will be useful in elucidating the molecular mechanisms that cause the strain specificity of algal symbiosis. RNAi methods have already been established for *Acanthamoeba*, and similar methods may apply to *M. viridis* (Lorenzo-Morales et al. 2005).

## Conclusion

We established an experimental system to collapse and re-establish photosymbiotic relationships between *M. viridis* and *Chlorella*. Our data showed that the photosymbiosis was not absolute but dispensable under the condition tested. Bleaching of *M. viridis* also made it possible to quantify the symbiotic abilities of each algal strain on certain criteria, which revealed that the symbiotic abilities of algae to *M. viridis* varied depending on the algal strains. This bleaching method will provide an important model tool not only for comparative analysis of hosts in the symbiotic and non-symbiotic states but also for observing the establishment process of photosymbiosis in live cell imaging.

## Acknowledgments

We thank Drs. Yukinori Nishigami, Yuu Ishii, and Masakado Kawata for technical suggestions; Dr. Keisuke Saito and the Cloud Dojo for Paper Writing Training program in the Photosynthesis Ubiquity project for supports in manuscript writing; Iwakuni City Microlife Museum for distribution of the *M. viridis* strain; Dr. Masashi Hayakawa for the suggestions on culturing the *M. viridis* strain. This work was supported by JST SPRING, Grant Number JPMJSP2108 and JSPS KAKENHI Grant Numbers 24KJ0740 (to D. Y.), 22H05668, 24H01462, 23H04962, 23H04957, 22H02697, 23K23960 (to S. Maruyama). Computational resources were provided by the Data Integration and Analysis Facility, National Institute for Basic Biology.

